# Machine Learning-Enhanced Mucosal Luminal Interface Metaproteomics Reveals Diagnostic Potential of Microbial Proteins in Pediatric Inflammatory Bowel Disease

**DOI:** 10.64898/2026.11.09.765240

**Authors:** Leila Ghanbari-Maman, Zahra Fekrirad, Anna Meyfour, Kaveh Kavousi

## Abstract

Inflammatory Bowel Disease (IBD), a chronic gastrointestinal disorder with two main subtypes, Crohn’s Disease (CD) and Ulcerative Colitis (UC), is associated greater diagnostic complexity in pediatric patients. Conventional diagnostic biomarkers, lack sufficient distinguishing power between IBD subtypes. This study aimed to investigate and compare proteomic and metaproteomic profiles of pediatric IBD (PIBD) patients and healthy controls to develop a diagnostic panel capable of accurately diagnose IBD and distinguish its subtypes.

We performed an in-depth evaluation of mucosal luminal interface samples from PIBD patients and healthy individuals, separately analyzing both the human proteome and microbial metaproteome data. The study involved selecting appropriate protein sets, employing feature selection techniques and machine learning across three intestinal regions: the terminal ileum, ascending colon, and descending colon.

Notably, panels composed solely of microbial biomarkers outperformed both human-only and combined human-microbial panels, achieving highly accurate results. Furthermore, the ascending colon emerged as the optimal sampling site.

This study uncovered disease relevant signaling pathways and delivered taxonomic as well as functional insights into the microbial dysbiosis of PIBD. A diagnostic panel built around microbial proteins—particularly those originating from the ascending colon—emerges as a promising tool for more precise stratification of PIBD subtypes.

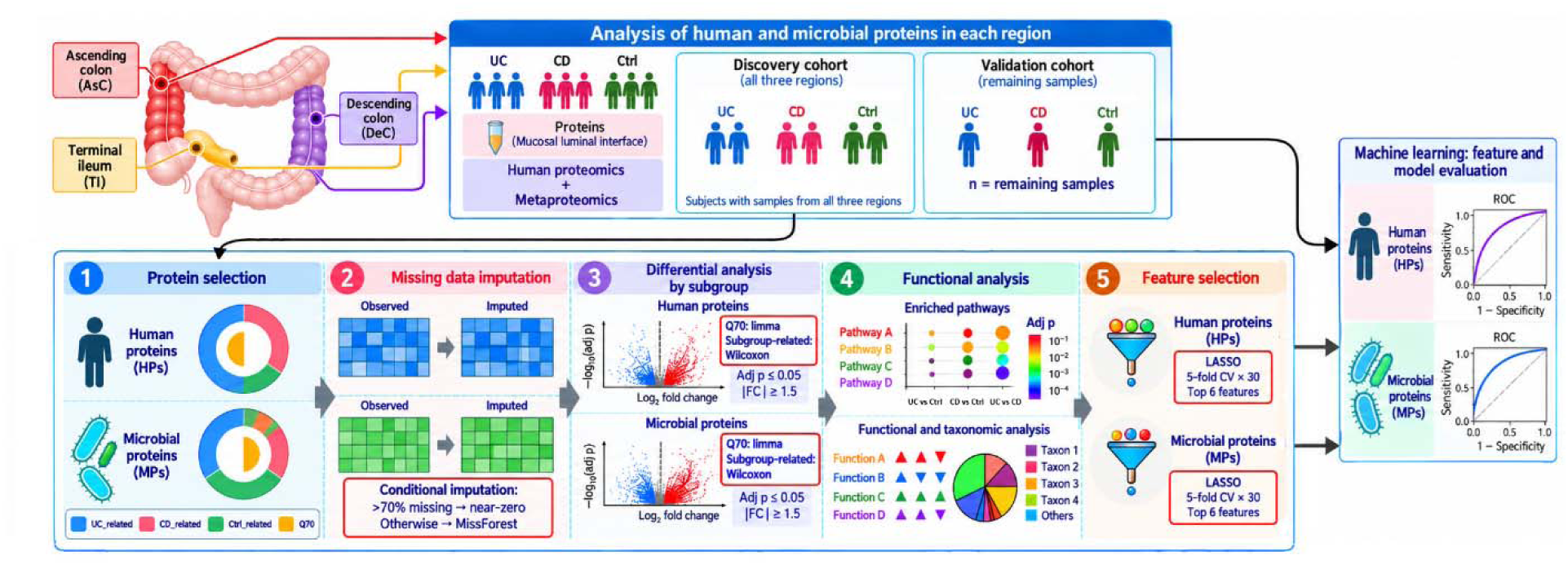

## 1. Introduction

Inflammatory Bowel Disease (IBD) is an autoimmune disorder, in which the immune system attacks the tissues of the gastrointestinal tract, leading to chronic inflammation. The IBD encompasses two subtypes: Crohn’s Disease (CD) and Ulcerative Colitis (UC). The global prevalence of IBD has surged in recent decades, becoming an emerging health concern Projections indicate that by 2030, more than 0.3% of the population in North America, Oceania, and parts of Europe will be affected, with Canada expected to reach approximately 1% [1, 2]. Notably, almost 25% of IBD patients are pediatric [3]. Research shows that Pediatric Inflammatory Bowel Disease (PIBD) exhibits distinct epidemiology, pathogenesis [6], and manifests as a more aggressive phenotype compared to adult IBD [4]. Additionally, the cost of care for pediatric CD is significantly higher than for adults, and UC costs are more than double those for adult cases [5]. Consequently, achieving new insights into the pathogenesis of PIBD and identifying specific signatures are crucial [6].

The number of studies conducted on pediatric cohorts is limited. Most of these investigations have mainly focused on finding expressional changes in human genes and introducing potential biomarkers in biopsy, serum, plasma, and stool samples of pediatric and adult IBD patients [7] [8], [9, 10] [11] [12]. However, these diagnostic biomarkers frequently lack the specificity and sensitivity necessary to distinguish CD from UC in pediatric population as well as PIBD patients from healthy controls.

To identify biomarkers with greater diagnostic power, it is essential to consider the molecular mechanisms involved in the pathogenesis of the disease. By focusing on key contributors to the disease process, diagnostic accuracy could be improved. The risk factors for IBD include genetics, smoking, hygiene, medications, microbiome, appendectomy, nutrition, stress, and physical activity [13]. Research into the impact of the gut microbiome on various diseases is continually growing, and it has been widely reported that the microbiome is closely linked to chronic illnesses [14]. There are several factors that can change the gut microbiota composition and function including host genetics, diet, age, lifestyle, hygiene, mode of birth, and medications like antibiotics [15]. The gut microbiome mainly consists of bacteria with a low percentage of archaea, eukaryotes, and viruses. The gut microbial gene profile was estimated to be 100–150-fold that of human genes with 99.1% originating from bacterial species [16]. Over 1,000 bacterial species live in the gastrointestinal tract, playing an important role in host homeostasis aspects like immunity, metabolism, nutrition, and defense against pathogens. Despite beneficial microbiota, which are immunosuppressive through the regulation of host immune cells, some harmful bacteria could stimulate inflammatory cytokines through the immune-cell interactions or induce intestinal damage [17]. The composition of the gut microbiota in IBD patients deviates from healthy individuals. It is characterized by reduced bacterial diversity and richness and an enrichment of opportunistic species. This microbial imbalance is called dysbiosis which co-occurs with transcriptional and metabolic changes in the gut [18]. Differences in microbial diversity have been demonstrated between IBD subtypes and healthy volunteers [19], thus microbial changes could serve as key diagnostic factors.

In the field of metagenomics, which mainly focuses on the microbial gene profiles, several studies have been conducted, and in some cases, taxonomy-based diagnostic panels have been proposed [20]. However, a more precise microbiome-based analysis is needed to understand the functional power of microbiome in diagnosis [21]. In this field, the use of protein-level data can provide valuable functional insights, as the pathogenesis of IBD is closely intertwined with microbial functions and changes throughout the illness, making it impossible to overlook the impact of the microbiome. However, studies utilizing proteomic and metaproteomic profiles in children with IBD remain scarce. Integrating proteomics and metaproteomics represents a novel approach to exploring the relationship between the host and microbiota in PIBD. Yan *et al.* conducted an analysis combining serum proteomics and stool metaproteomics to investigate the impact of obesity on PIBD [22]. Zhang *et al.* provided proteomic and metaproteomic data to find out the associations between the microbiome and intestinal extracellular vesicles in PIBD patients [23]. These findings would provide a clearer understanding of the microbiome’s impact on disease pathogenesis, particularly its potential as a key diagnostic biomarker for IBD. In comparison with the profiles of fecal microbiota, it has been found that gut mucosa-adhering microbiota can better differentiate IBD patients from controls by revealing some noteworthy biomarkers [24]. In this study, the human proteome and metaproteome profiles of PIBD were analyzed to identify an accurate set of biomarkers for PIBD diagnosis by using a dataset comprising mucosal luminal interface samples of 71 patients from three gut regions. The analysis pipeline was separately conducted for human proteome and microbial metaproteome profiles of Terminal Ileum (TI), Ascending Colon (AsC), and Descending Colon (DeC) regions. We carefully selected a specific protein set with strong discriminative potential and used an imputation method that maintains the minimal error in the intensity of each protein within the two subgroups of IBD (UC and CD) as well as healthy controls. Separately analysis of proteome and metaproteome profiles to identify differentially expressed proteins followed by enrichment analysis led to find relevant pathways to IBD. Furthermore, a machine learning pipeline was applied to the proteomic and metaproteomic data to find the best diagnostic panel for PIBD in each region. Additionally, we attempted to combine host- and microbe-based features to check whether smaller-sized biomarker panel could be identified for the three regions of gastrointestinal tract. Using mucosal luminal interface (MLI) samples, we discovered microbe-based panels in AsC region that could predict the patients’ condition with 100% accuracy.

## 2. Methods

### 2.1. Study population

In this study, we utilized a dataset (PXD007819 and PXD007959) comprising 176 MLI samples collected from three gut regions: 62, 58, and 56 samples from the AsC, DeC, and TI, respectively. These samples were extracted from a group of 71 pediatric individuals including 25 CD, 22 UC treatment-naïve patients, and 24 non-IBD control subjects. All IBD patients were in flare mode (Activity index>15) (Supplementary Table S1). There were no significant differences in age among healthy control, CD, or UC patients, nor in the number of females and males in each subgroup. Notably, samples from 42 patients who had available samples from all three gut regions were used for biomarker discovery, and the remaining samples from other individuals were used to validate the candidate panels (Table 1).

**Table 1.** Cohort characteristics of the study.

|  | Ctrl (n=24) | CD (n=25) | UC (n=22) |
|---|---|---|---|
| Gender (Male, Female) | 10 , 14 | 16 , 9 | 11 , 11 |
| Median Age | 15.0 | 13.8 | 14.9 |
| Inflammatory Status |  |  |  |
| TI (inf, non-inf) | 0 , 21 | 16 , 3 | 0 , 16 |
| AsC (inf, non-inf) | 0 , 22 | 11 , 11 | 12 , 6 |
| DeC (inf, non-inf) | 0 , 20 | 11 , 9 | 17 , 1 |

TI: Terminal Ileum, AsC: Ascending Colon, DeC: Descending Colon, Ctrl: Healthy control, CD: Crohn’s Disease, UC: Ulcerative Colitis, Inf: Inflamed and non-inf: non-inflamed.

### 2.2. Preprocessing

The samples successfully passed quality control. To achieve the most accurate identification, mass spectrometry analysis was performed using iMetalab [25] and the built-in MaxQuant software on all 176 MS files and 8 samples reported in [23] as the quality control samples in a single run. The analysis was conducted against the previously introduced database [23] according to reported values for each parameter. Proteins that were contaminants or mapped to the reverse sequence of a protein in the decoy database were removed. Proteins with fewer than 2 unique peptides were also excluded. For further analysis, we utilized the DEP package (version 1.20.0) in R software [26]. Normalization of intensities was performed using the VSN method on summarized experiments. In order to identify biomarkers during the discovery phase, we separated data obtained from the 42 specific patients who had samples from all three regions, while the remaining patients were used as an independent test set for validation in each region. The analysis pipeline for each region on these samples was set up as follows.

### 2.3. Intestinal region-based analysis

#### Protein filtration

In the first step, we pruned the proteins based on several filters. The proteins not identified in the samples of the region being analyzed were removed and then the missing rate of each protein was separately calculated for the three sample subgroups: CD patients, UC patients, and healthy control. Since the primary goal of biomarker discovery is to identify subgroup-specific markers that are not present in the others, two classes of proteins were selected for further analysis: subgroup-related proteins and Q70 proteins. Subgroup-related proteins were those quantified in at least 70% of one subgroup while being found in less than 30% of at least one of the other subgroups, termed CD-related, UC-related, and Ctrl-related proteins. The union of all these proteins is referred to as subgroup-related proteins. Furthermore, identified proteins in at least 70% of each subgroup were separated as Q70 proteins. We applied Q70 and subgroup-related proteins as the filtered proteins for further analysis. By applying such filters, only bacterial and human proteins remained, while viral, fungal, and other proteins were entirely removed.

#### Missing data imputation

According to the definition of the subgroup-related proteins, they do not exhibit a similar distribution pattern of missing values in the subgroups and they are characterized by having Missing Not At Random (MNAR) intensity values. Since the accuracy of imputation is primarily influenced by the MNAR rate rather than other missingness rates, and the choice of imputation method can significantly impact downstream analysis, it is crucial to select an appropriate imputation method for the filtered proteins.

Jin *et al.* compared the most popular and common imputation methods used for the proteomics data with different rates of missingness in the intensities. They proposed the Random Forest (RF) imputation method as the most accurate for the proteomics data, due to the lowest error rate, presenting the best relevant pathways and a high number of true positives [27]. As our strategy for the protein selection was based on finding the MNAR proteins (subgroup-related proteins) as well as the proteins whose expression was detected in most samples (Q70 proteins), we employed a conditional imputation method. For the proteins with more than 70% missingness in a subgroup’s samples, we imputed missing values by a random value close to zero (normal distribution by mean=1, sd = 1), while the Random Forest method in MissForest package of R was applied for the other proteins [28].

#### Statistical analysis

Since the t-test demonstrated a significant difference in mean log intensity between the Human Proteins (HPs) and Microbial Proteins (MPs) in each region, the downstream analysis was performed separately. By using the limma method on Q70 proteins, the proteins whose expression showed significant changes (adjusted *p*-value <= 0.05, absolute fold change >= 1.5) in at least one of the comparisons between two subgroups, were identified as significant proteins. For the subgroup-related proteins, the Wilcoxon test was used to find proteins with different intensity distributions in subgroups.

#### Enrichment analysis of HPs and functional and taxonomical analysis of MPs

EggNOG-mapper [29] was used for taxonomical and functional profiling of significant MPs. By grouping the significant MPs based on the Cluster of Orthologues Groups (COGs), the most abundant COGs were identified in each comparison of the subgroups. Taxonomic profiling of significant MPs was performed at the family level. The pathway enrichment analysis of significant HPs was done by EnrichR to identify the most important pathways in subgroup comparisons.

#### Feature selection

By using the Least Absolute Shrinkage and Selection Operator (LASSO) method in R [30], multinomial feature selection was separately performed for significant HPs and MPs. To account for randomness, we repeated 5-fold cross validation of LASSO 30 times and calculated the average coefficient for each protein. The top 6 HPs and MPs with the highest average coefficients were selected as potential features for further evaluation.

#### Biomarker discovery and validation

During the machine learning phase, the Random Forest algorithm was applied to various combinations of features, namely 2, 3, 4, 5, and 6 markers. We separately evaluated 6 human features and 6 microbial features to determine the maximum power of the HP- and MP-based biomarker panels in discriminating the subgroups. Alongside, we tried to achieve the most accurate possibly combined panels with the minimum number of biomarkers. The process of learning was performed with 5-fold cross-validation on the training set, which encompassed 70% of the samples from the discovery cohort including 42 patients. Subsequently, the model was tested on the remaining 30% of the discovery cohort and the mean accuracy was reported as the training accuracy. For each combination, the accuracy and F1-score of the learned machine were also calculated on a separate unseen validation cohort. This cohort included 14, 20, and 16 samples for the TI, AsC, and DeC regions, respectively. The panels which had the most accuracy on the validation cohort were reported as the diagnostic biomarker panels. The average accuracy and F1-score were calculated for both HP- and MP-based panels as well as combined panels on the validation set, and the AUC of each panel in each subgroup versus the others was also reported.

## 3. Results and discussion

### 3.1 Preprocessing, Filtering and identification of proteins

A total of 53,694 protein groups were identified and quantified using iMetalab and the built-in MaxQuant, including 48,307 MPs, 3,344 HPs, and 2043 other proteins from various microorganisms (viruses, fungi, etc.). Within this set, 480 proteins were either mapped to the decoy database or considered contaminants. Following the exclusion of these proteins and those with less than 2 unique peptides, 22,995 proteins remained.

To evaluate the diagnostic potential of each region and compare biomarker candidates across intestinal regions, we utilized samples from 42 patients who had samples from all three regions (Figure 1A). The proteins identified in these 42 individuals were employed as the discovery set, while the remaining samples were reserved for the validation test set. Figure 1B illustrates the number of samples for each region as well as the inflammation information for the subgroups in the discovery set.

**Figure 1.**
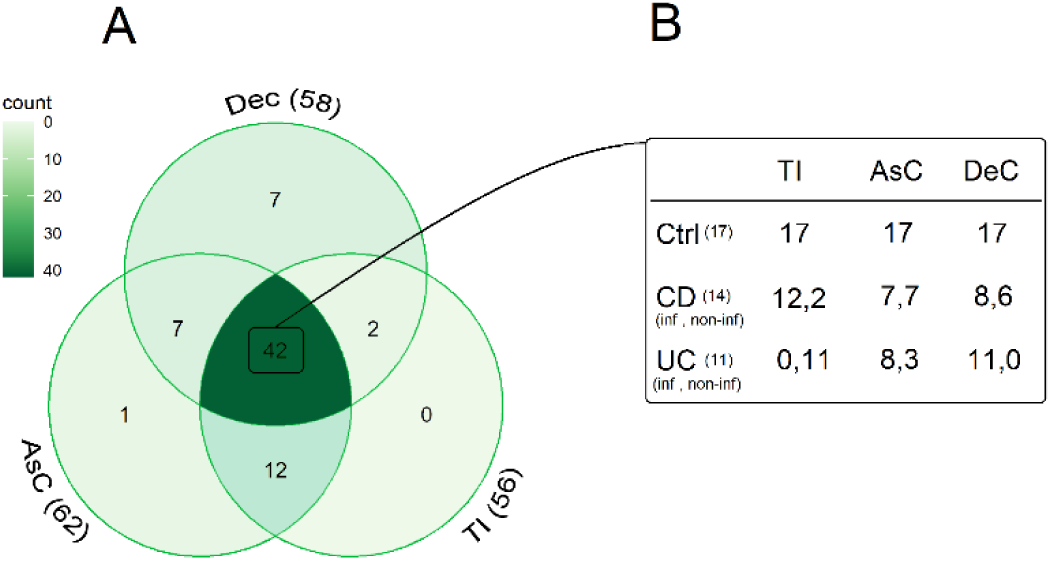
42 patients had samples for three regions, which used as a discovery cohort. **(A)** Venn diagram shows the distribution of samples in the TI, AsC, and DeC. The 42 patients with samples from all three regions were used as the discovery cohort and the rest as the validation cohort, **(B)** Detailed information about subgroups and their inflammatory status. TI: Terminal Ileum, AsC: Ascending Colon, DeC: Descending Colon, UC: Ulcerative Colitis, CD: Crohn’s Disease, inf: inflamed, and non-inf: non-inflamed, Ctrl: Healthy control.

Among the 22,995 proteins, 18,908 (16,456 microbial, 2,452 human) proteins were detected in TI, while 17,314 (14,942 microbial, 2,372 human) and 18,319 (15,887 microbial, 2,432 human) proteins were found in AsC and DeC, respectively. These differences in the number of MPs may reflect the heterogeneity of microbial population in different parts of colon, consistent with previous studies which have demonstrated the diversity of the microbiota communities in different regions of the gut [31, 32]. It may have resulted from underlying differences in gut physiology, host effects on the microbiome, or the various roles of gut regions in food digestion, absorption, and fermentation [33–35].

After filtration, two groups of identified proteins were selected for further analysis: the subgroup-related proteins that were quantified in at least 70% of samples in a subgroup while being expressed in less than 30% of at least one of the other two subgroups, and Q70 proteins whose expression was observed in >70% of the samples from all three subgroups. By applying these filters, only human and bacterial proteins remained, while proteins from other microorganisms failed to meet sufficient criteria for further analysis. Consequently, in total, 736 (373 microbial and 363 human), 980 (526 microbial and 454 human), and 808 (428 microbial and 380 human) filtered proteins were identified in TI, AsC, and DeC, respectively. Divergent conditions throughout the gastrointestinal tract led to regional differences in the microbial communities of different parts. As a result, the microbial composition varies between colonic regions with distinct microenvironments [36]. The highest number of filtered MPs and HPs were detected in AsC region, which reflects that AsC may undergo more changes than the other regions.

With an in-depth look at the expression pattern of subgroup-related proteins, certain proteins were identified that could not be detected in other subgroups at all, termed extreme MNAR proteins. These proteins can provide essential information for understanding differences in pathogenesis and distinguishing different subgroups. In the context of TI, just one extreme MNAR microbial protein was found which was not identified in UC samples. Notably, in the DeC region, no extreme MNAR proteins were found as all proteins were detected in at least one sample of each subgroup. In the AsC region, there were 46 extreme MNAR proteins including three HPs which were not detected in the CD patients (G-protein subunit γ 5 (GNG5), Heterogeneous Nuclear RiboNucleoProtein C (HNRNPC), and Myristoylated Alanine Rich protein Kinase C Substrate (MARCKS)) (Figure 2). Furthermore, 28 and 13 MPs were not observed in the UC and healthy control subgroups, respectively. As shown in Figure 2, both UC and healthy control samples lacked 2 MPs.

**Figure 2.**
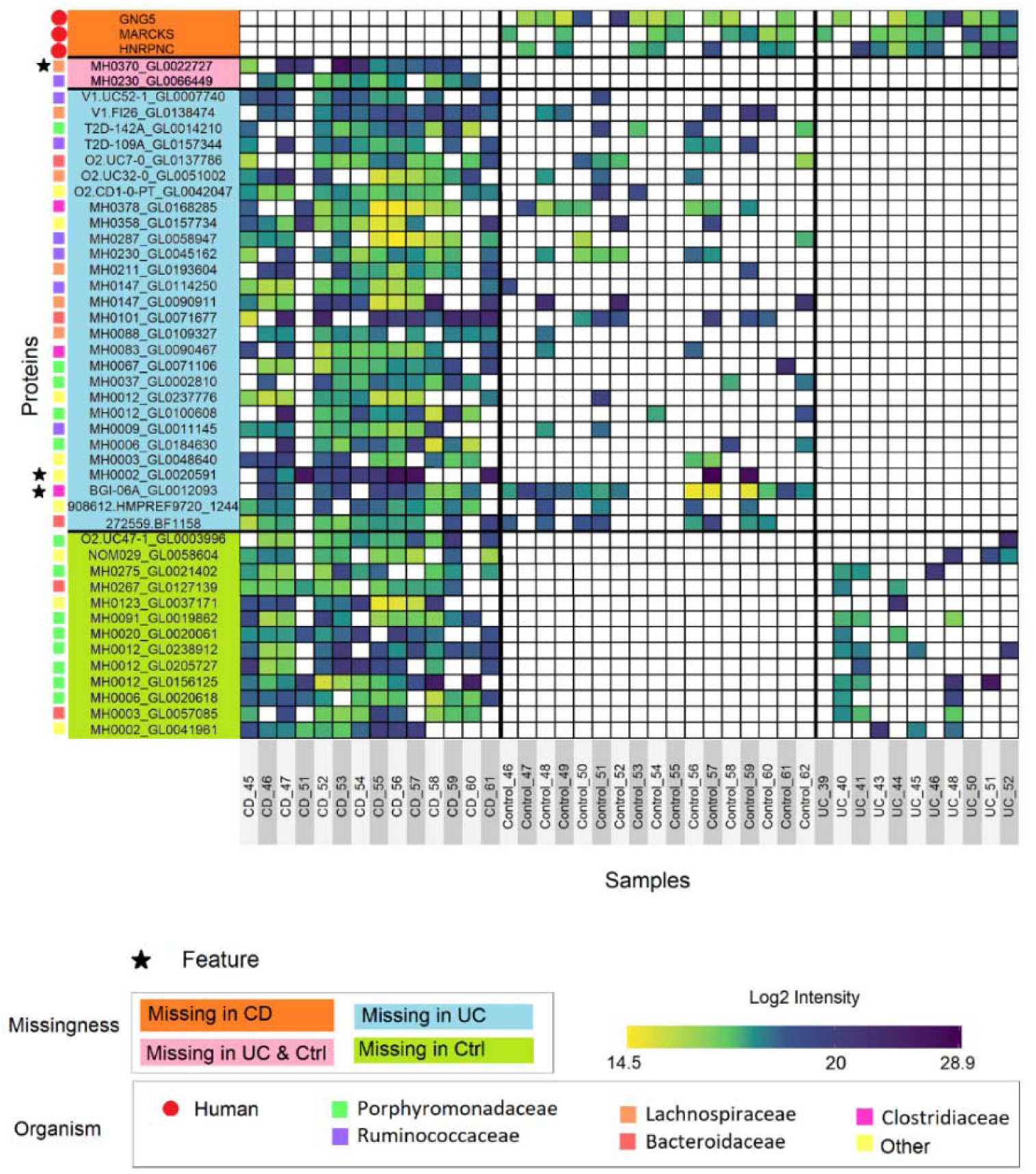
Ascending Colon with the most extreme Missing Not At Random MicrobialProteins. The intensity heatmap of proteins that were not detected in the samples of at least one subgroup. The proteins are shown in 4 categories: not detected in CD subgroup (Orange), not detected in the Ctrl and UC subgroups (Pink), not detected in UC subgroup (Blue), and not detected in the Ctrl subgroup (Green). Each star represents a feature to be selected for machine learning through subsequent analysis. Circles indicate human proteins. The taxonomies to which each proteins belongs are indicated by the colors next to the heatmap diagram. UC: Ulcerative Colitis, CD: Crohn’s Disease, Ctrl: Healthy control.

The taxonomies with the most abundant MNAR proteins in AsC were *Porphyromonadaceae* and *Bacteroidaceae* families with 13 and 5 proteins respectively, both from phyum *Bacteroidota.* Furthermore, 7, 6 and 3 MNAR proteins belonging to *Ruminococcaceae*, *Lachnospiraceae* and *Clostridiaceae* families from phylum *Bacillota*, respectively. The impact of these taxonomies on IBD, has been reported before [37–39]. The *Bacillota*/*Bacteroidota* ratio (formerly *Firmicutes*/*Bacteroidetes* (F/B)) has emerged as a potential biomarker for health. An increased ratio has been observed in gastrointestinal conditions such as IBD and irritable bowel syndrome (IBS). However, the microbiome’s complexity could limit the reliability of this health indicator [40].

It seems that most changes in microbial population occurred in AsC, as this region exhibited the highest number of extreme MNAR proteins. These observations strengthen the hypothesis that MPs of AsC may be more suitable candidates for discovering IBD diagnostic biomarkers. Among the extreme MNAR HPs in AsC, GNG5 has previously been reported as a biomarker for Melanosis Coli, which is associated with the pigmentation of the colon [41]. Numerous studies supported the notion that pigment deposition is correlated with epithelial cell apoptosis. It is now recognized that any condition inducing mucosal cell apoptosis, such as IBD, can lead to this pigmentation [42]. Another HP extreme MNAR protein was HNRNPC, a regulator/reader of m6A whose methylation has been implicated in UC pathogenesis [31]. Targeting m6A is considered a potential adjuvant therapy for IBD [32].

MARCKS, another extreme MNAR human protein, contributes to regulating toll-like receptor 4 signaling in macrophages. CRISPR-CAS9-mediated knockout of MARCKS in macrophages downregulated the production of TNF-α and IL-6. It can contribute to specific biological processes such as innate immune responses, inflammatory responses, and cytokine production. MARCKS appears to be a key regulator of inflammation, and its inhibition might hold therapeutic potential for inflammatory diseases [43–45].

### 3.2 The landscape of protein alteration

#### The microbial protein profiles were analyzed separately from the human proteomes

Missing proteins with more than 70% missingness in a subgroup were imputed with a random number close to zero, while other proteins were imputed using the RF method. A t-test was conducted to compare the mean intensity of MPs and HPs, yielding a *p*-value < 2.2 e-16, which indicates a significant difference between the frequency of protein expression in the two groups. The confidence intervals were calculated as [2.4 to 2.7], [7.2 to 7.6] and [2.0 to 2.3] for TI, AsC, and DeC, respectively. These results indicate that the log2 intensity of MPs is significantly lower than that of HPs, consistent with findings reported by Zhang *et al.* [23]. This difference was most prominent in the protein profiles of AsC region. Thus, it is recommended to perform separate microbial and human proteome analyses to identify differentially expressed proteins and gain more accurate insights.

The test-diff function within the DEP package, integrating the limma function, was used to identify significant differentially expressed HPs and MPs among the Q70 proteins (adjusted *p*-value <= 0.05 and absolute fold change >= 1.5 in at least one comparison between two subgroups). For the subgroup-related proteins, Wilcoxon’s Rank Sum test was used to identify proteins with distinct intensity distribution patterns. We found 49, 411, and 348 significant HPs, and 351, 399, and 113 significant MPs in the TI, AsC, and DeC regions, respectively (Supplementary Table S2). Some proteins exhibited differential expression patterns between two subgroups, while others were significant across all three subgroups.

Figure 3 illustrates the common and distinct significant MPs and HPs among CD, UC, and healthy control subgroups across the three different gut regions. In the TI region, the number of significant MPs exceeded that of HPs (Figure 3A, D), the most significant proteins were found in UC patients compared to healthy controls (Figure 3D) while in the DeC region, this pattern was reversed, with HPs being more dominant (Figure 3C, F). However, in both these regions, in contrast, the AsC region demonstrated a balance in terms of the number of significant MPs and HPs (Figure 3B, E), while the most pronounced differences in both HPs and MPs were observed when comparing IBD subtypes (Figure 3B, E). Hence, as shown in Figure 3G, the most changes in the human proteome profiles were identified in the AsC and DeC regions, while as depicted in Figure 3H, the microbial profiles of TI and AsC regions underwent greater changes (Figure 3H). Considering this, it appears that the most dysregulation of HPs and MPs could be seen in the AsC region of PIBD patients.

**Figure 3.**
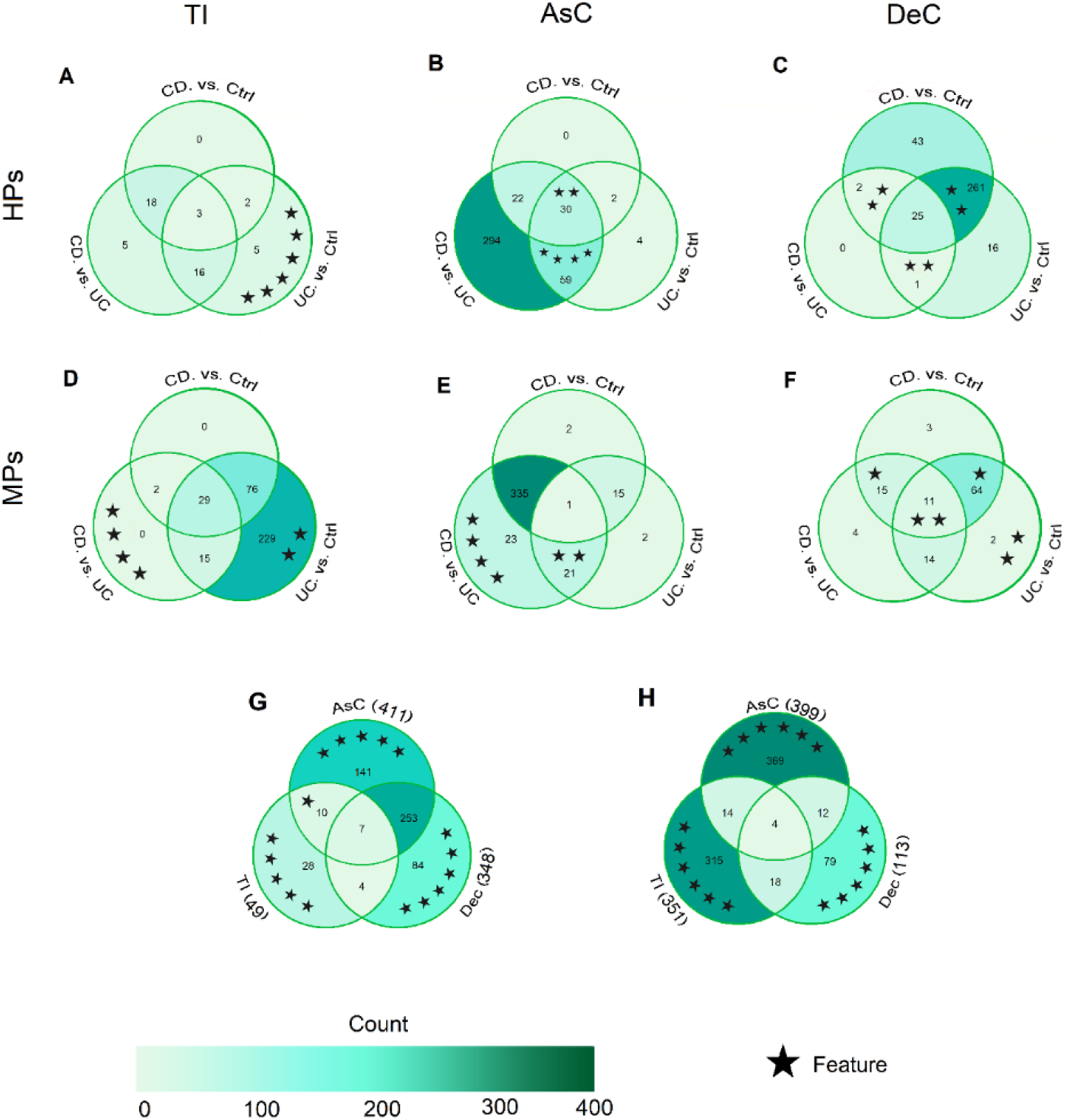
The landscape of significant HPs and MPs shows more alterations in the MLI samples of AsC compared to other regions of PIBD patients. Venn diagram of the significant HPs **(A, B, C)** and MPs **(D, E, F)** in comparisons of TI, AsC, and DeC proteomes and metaproteomes of CD, UC, and healthy control individuals. Venn diagram of the common and distinct significant HPs **(G)** and MPs **(H)** among the three gut regions. The significant proteins were the union of significant subgroup-related and the Q70 proteins with absolute fold change >=1.5 and adjusted p-value <=0.05. Each star represents a feature to be selected for machine learning through subsequent analysis. The numbers in parentheses (G, H) indicate the total number of significant proteins identified for each intestinal region. HPs: Human Proteins, MPs: Microbial Proteins, TI: Terminal Ileum, AsC: Ascending Colon, DeC: Descending Colon, UC: Ulcerative Colitis, CD: Crohn’s Disease, Ctrl: Healthy control.

Venn diagrams (Figure 4) illustrate the significant proteins in the three gut regions based on the two mentioned criteria for filtration, from another perspective. The significant Q70 MPs were observed exclusively in the TI region, indicating that microbial profile changes in this area are mainly limited to variations in expression rather than loss or new expression of microbial proteins. Regarding HPs, the changes are not noticeable, suggesting that the disease has not yet caused severe changes in the human proteome profile of this region. Unlike the TI region, where the number of significant Q70 human proteins was low (10), 308 and 316 HPs were differentially expressed in AsC and DeC, respectively.

**Figure 4.**
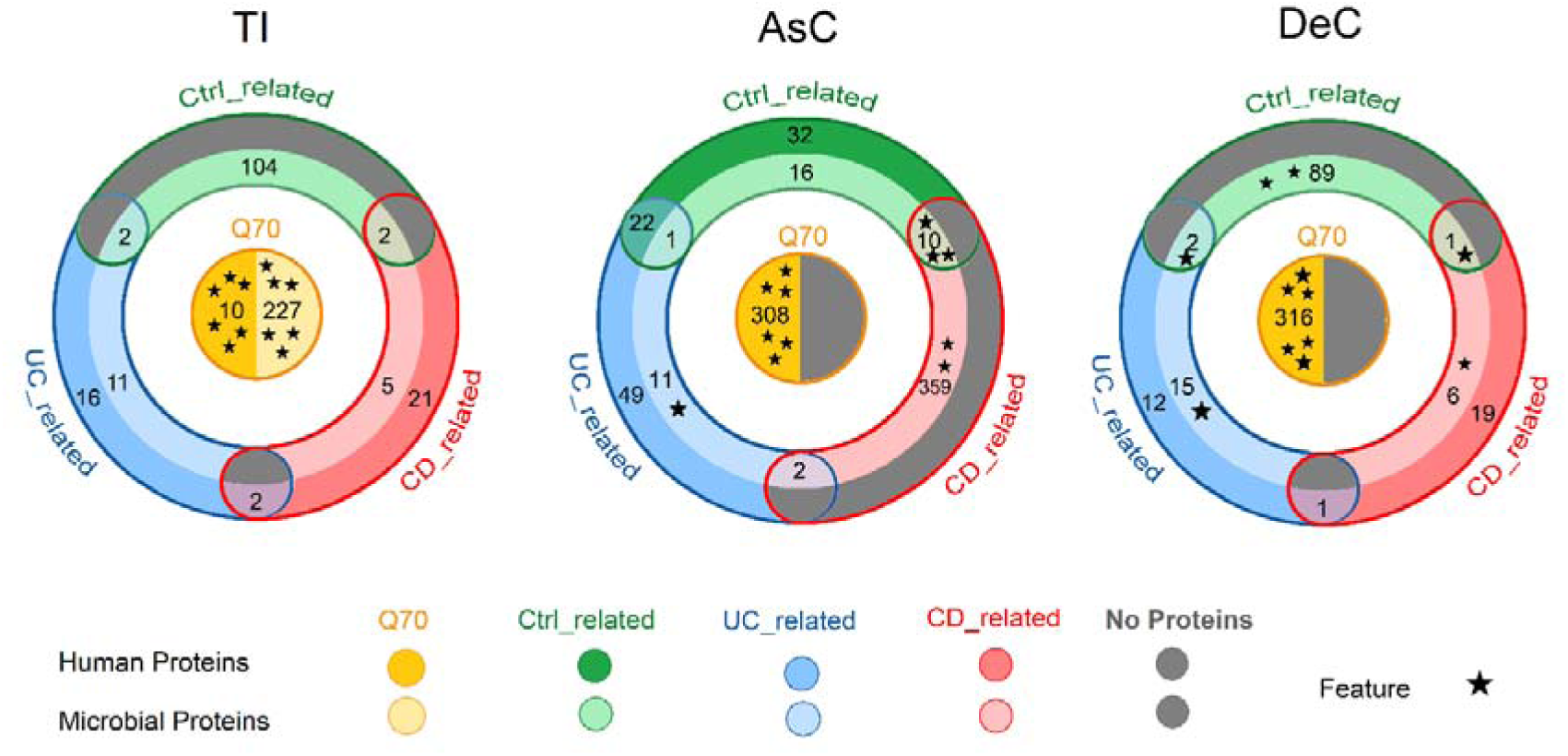
Absent or new expression of microbial proteins in MLI samples of PIBD. Venn diagram of the significant human and microbial proteins based on the number of CD-related, UC-related, and Ctrl-related proteins as well as Q70 proteins with absolute fold change>=1.5 in TI, AsC, and DeC. Each star represents a feature to be selected for machine learning through subsequent analysis. TI: Terminal Ileum, AsC: Ascending Colon, DeC: Descending Colon, UC: Ulcerative Colitis, CD: Crohn’s Disease, Ctrl: Healthy control.

In each region (Figure 4), it is also evident that no significant subgroup-related HPs were found in a condition with the maximum number of significant subgroup-related MPs, such as Ctrl-related proteins in TI and DeC as well as CD-related proteins in AsC. Conversely, when the number of significant subgroup-related HPs in a region reaches its maximum, changes in the subgroup-related MPs are at their lowest, such as CD-related proteins in TI and DeC regions, as well as UC-related proteins in AsC. It is worth noting that the most significant HPs were Q70 proteins whose expression was altered in the MLI samples of PIBD patients. Some of these proteins have been identified as diagnostic biomarkers in previous studies [46, 47]. However, especially in the colonic region, we primarily observed the removal or presence of new MPs, whose important and undeniable contribution should be considered in the pathogenesis and treatment of this disease.

The significant proteins in each region were used for correlation evaluation and Principal Component Analysis (PCA), revealing distinct separations among subgroups (Supplementary FigureS1). The correlation heatmaps and PCA plots demonstrated that the microbiome’s significant proteins (Supplementary Figure S1D-F and J-L) exhibited better discriminatory power compared to the human proteins (Supplementary Figure S1A-C and G-I) in the respective regions.

Intriguingly, the discrete metaproteome profiles of all regions indicated a clear potential to separate the three subgroups (Supplementary Figure S1D-F & J-L), with AsC showing the most distinct segregation (Supplementary Figure S1E and K). Despite all UC samples in TI being from a non-inflammatory region, a noticeable separation was observed among the subgroups in both significant MPs and HPs (Supplementary Figure S1G and J). This result, along with a large number of significant MPs observed between the UC and Ctrl subgroups in the TI region (Figure 3D), is consistent with the previous findings that UC patients showed ileal barrier depletion and a decrease in the microbiome diversity of their ileum even in the absence of inflammation [48].

Several HPs we identified as significant proteins have been also reported in other studies. In particular, B2M, ECH1, HADHB, and TF, which we found as significant proteins in the AsC region, have been also previously suggested by Starr *et al.* as potential biomarkers for distinguishing CD from UC samples, using the proteome analysis of AsC biopsies [8]. It seems that these proteins are secreted from the biopsy to the lumen. Three other significant HPs, CAT, TKT, and ANXA3 have been previously proposed as part of a biomarker panel for discriminating UC from Ctrl using MLI samples from AsC and DeC regions of PIBD [46].

Moreover, twenty-one proteins (A2M, ANXA1, ANXA5, AZU1, CORO1A, CPNE3, ELANE, HBA1, HBB, HP, LCP1, LTF, MMP8, MPO, PRTN3, RETN, S100A12, S100A8, S100A9, S100P, and SERPINB1) reported as biomarkers for diagnosing PIBD flares in stool samples [12], also exhibited significant differentially expression in our study. LCN2, reported before as a biomarker for the disease activity of UC patients [37], was also found to be significant in our study. SLC26A2 whose intensity significantly altered among subgroups in TI has previously been reported as a biomarker for discriminating CD, UC, and IBS [49], in our study. Also, we have 183 common significant proteins with the IBD-related gene list reported by Alfonso Perez *et.al.* [50]. It appears that the methodology employed in this study yields reliable results that are not dependent on the dataset and may have similar outcomes when applied to other datasets as well.

### 3.3 Functional enrichment of host and microbiome proteome profiles in PIBD

#### Significant subgroup-related HPs enhance pathway enrichment

For the pathway enrichment analysis, we employed EnrichR based on the MSigDB Hallmark 2020. Highlighting proteins enriched in at least one pathway with an adjusted p-value of <= 0.05, Figure 5 effectively demonstrates their significance (details in Supplementary Table S3).

**Figure 5.**
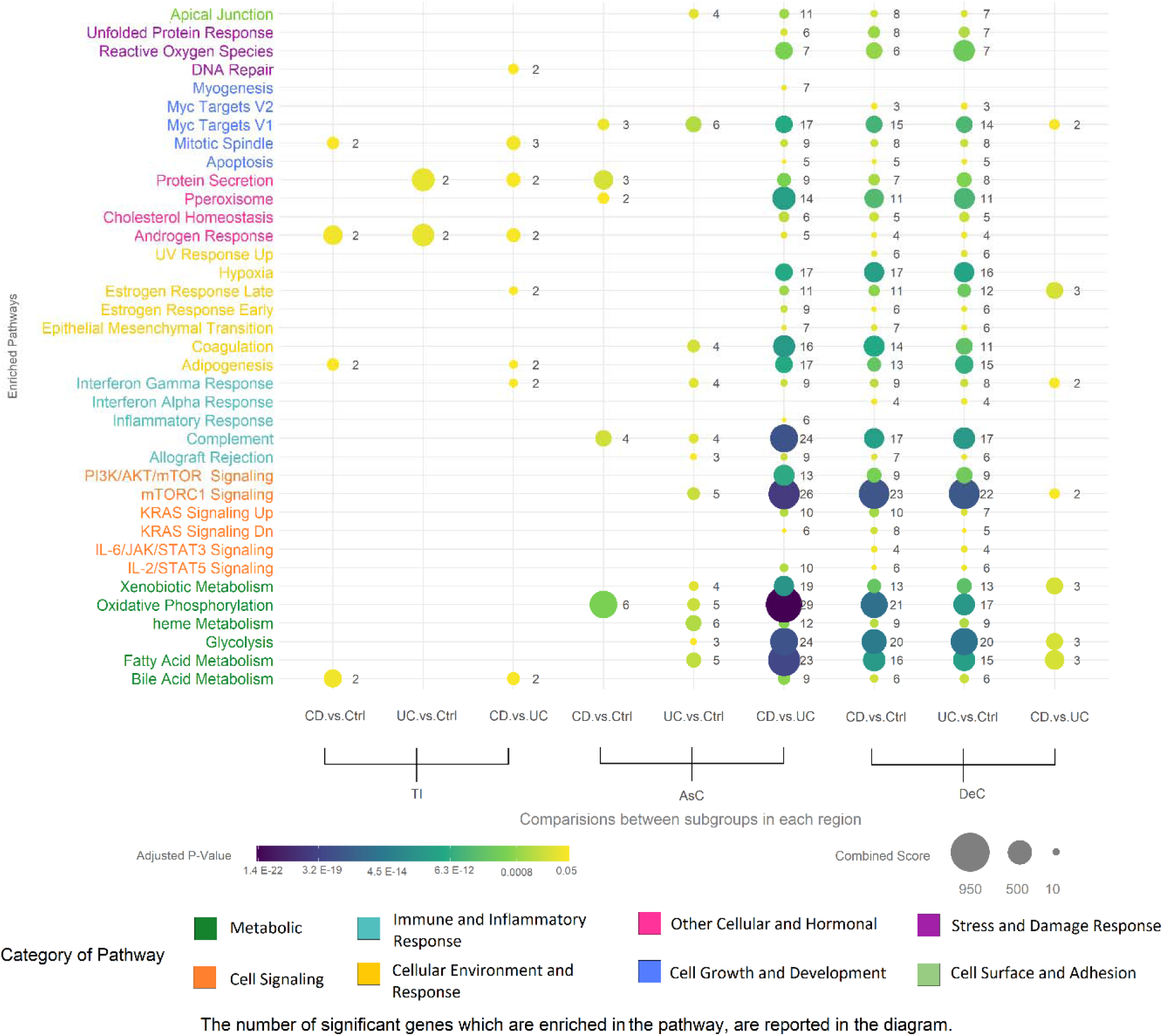
In comparing IBD subtypes, the most significant changes in signaling pathways are observed in the AsC region. Host significant pathways (adjusted *p*-value <=0.05) which were enriched in three regions TI, AsC, and DeC among three subgroups UC, CD, and Control. The size and color of the bubbles show the score and adjusted *p*-value of the pathways, respectively. The number of gene member for each pathway is reported next to the corresponding bubbles. TI: Terminal Ileum, AsC: Ascending Colon, DeC: Descending Colon, UC: Ulcerative Colitis, CD: Crohn’s Disease, Ctrl: Healthy control.

In a comprehensive analysis of the enrichment results, similar signaling pathways were found to be enriched in the DeC region when both IBD subtypes were compared with healthy individuals. Given the substantial overlap of significant proteins between DeC and AsC regions (Figure 3G), this enrichment pattern was also observed in the AsC region, with the difference that firstly, a large number of significant proteins were obtained by comparing the IBD subtypes and secondly, pathways enriched in the AsC region had higher significance scores and a greater number of related genes. Notably, the TI region exhibited the least enriched pathways compared to the other two regions, which may be attributed to the limited number of significant HPs in this region (Figure 3G). Furthermore, significant enriched pathways across different regions revealed that the disparities between IBD subtypes in the AsC are notably more pronounced than those observed in the DeC and TI, as clearly depicted in the scales of PCA Supplementary Figure S1G-I.

Several pivotal signaling and metabolic pathways were identified that play crucial roles in the pathogenesis of the disease. Among them, considering gene ratios, adjusted p-values, and combined scores, Oxidative Phosphorylation emerged as a cornerstone. This pathway was enriched in the DeC region of IBD patients and particularly in the AsC region of CD patients when compared to UC ones. This observation resonates with prior studies documenting reductions in specific complexes of Oxidative Phosphorylation in IBD, notably in UC [51]. This finding underscores the centrality of mitochondrial dysfunction and energy production deficiency in IBD. Mitochondrial dysfunction has been associated with the increased production of reactive oxygen species (ROS), another enriched pathway in the Stress and Damage Response category, leading to oxidative stress, which is a known contributor to the inflammatory characteristic of IBD [52].

The enrichment of the Glycolysis and Fatty Acid Metabolism pathways, especially in AsC and DeC, highlights their important role in IBD pathogenesis, which is consistent with previous research [53, 54] Glycolysis affects the gut microbiota, mucosal barrier function, and immune system modulation. Dysregulation of pathways like Glycolysis and Fatty Acid Metabolism plays a critical role in metabolic reprogramming of immune cells [54], where a shift from oxidative phosphorylation to glycolysis, known as the Warburg effect, supports the energy demands of proliferating inflammatory cells [55]. mTORC1 Signaling and PI3K/AKT/mTOR Signaling were also significantly enriched in IBD, indicating that dysregulation of cell growth, proliferation, and survival contributes to the disease pathogenesis. The mTOR pathway which is well-known for its role in regulating T cell differentiation and function, has been reported to promote UC via COX-2-mediated Th17 responses [56]. Hyperactivation of this pathway can result in excessive immune responses that exacerbate intestinal inflammation. This pathway is also connected to glycolysis and oxidative phosphorylation [57], showing the complex relationship between energy metabolism and cell signaling in IBD pathogenesis. It presents a promising approach for potential therapeutic intervention in IBD [58].

Bile Acid Metabolism was also significantly affected, with evidence pointing to dysregulation in bile acid synthesis and recycling, which is known to influence gut microbiota composition and intestinal inflammation [59]. Bile Acids produced by the liver, are crucial for the digestion and absorption of lipids and liposoluble vitamins in the intestine. Hepatocytes excrete conjugated bile acids with either taurine or glycine into the bile duct. When reaching the intestine, particularly the colon, they were metabolized by the gut microbiota. In fact, Bile Acids impact the composition and activity of numerous intestinal bacteria, promoting the growth of certain species and suppressing others. They also play a key role in controlling the synthesis of antimicrobial peptides in the intestine and are essential for maintaining a healthy and intact intestinal barrier [60].

As an attractive point, alterations in Heme Metabolism have been also observed in IBD patients, which is consistent with previous findings [61, 62]. Choby *et al.* demonstrated the importance of the iron-containing cofactor heme for bacterial pathogens in the disease development [63]. Likewise, iron exerts a significant impact on the composition of the gut microbiota [64]. It is worth noting that anemia stands out as one of the most common manifestations of IBD, particularly in pediatric patients [65].

#### AsC is different from the other regions in the functional analysis of the significant MPs

To delve into the functional characteristics of microbial composition, we employed eggNOG-mapper. Figure 6 clearly illustrates the Cluster of Orthologue Groups (COGs) of significant MPs, with the size of the triangles reflecting the mean intensity fold change of corresponding MPs. The number of significant MPs for each COG is also displayed in the respective triangle. Up-pointing and down-pointing triangles show positive and negative intensity fold changes when comparing one subgroup to another, respectively (additional details in Supplementary Table S4).

**Figure 6.**
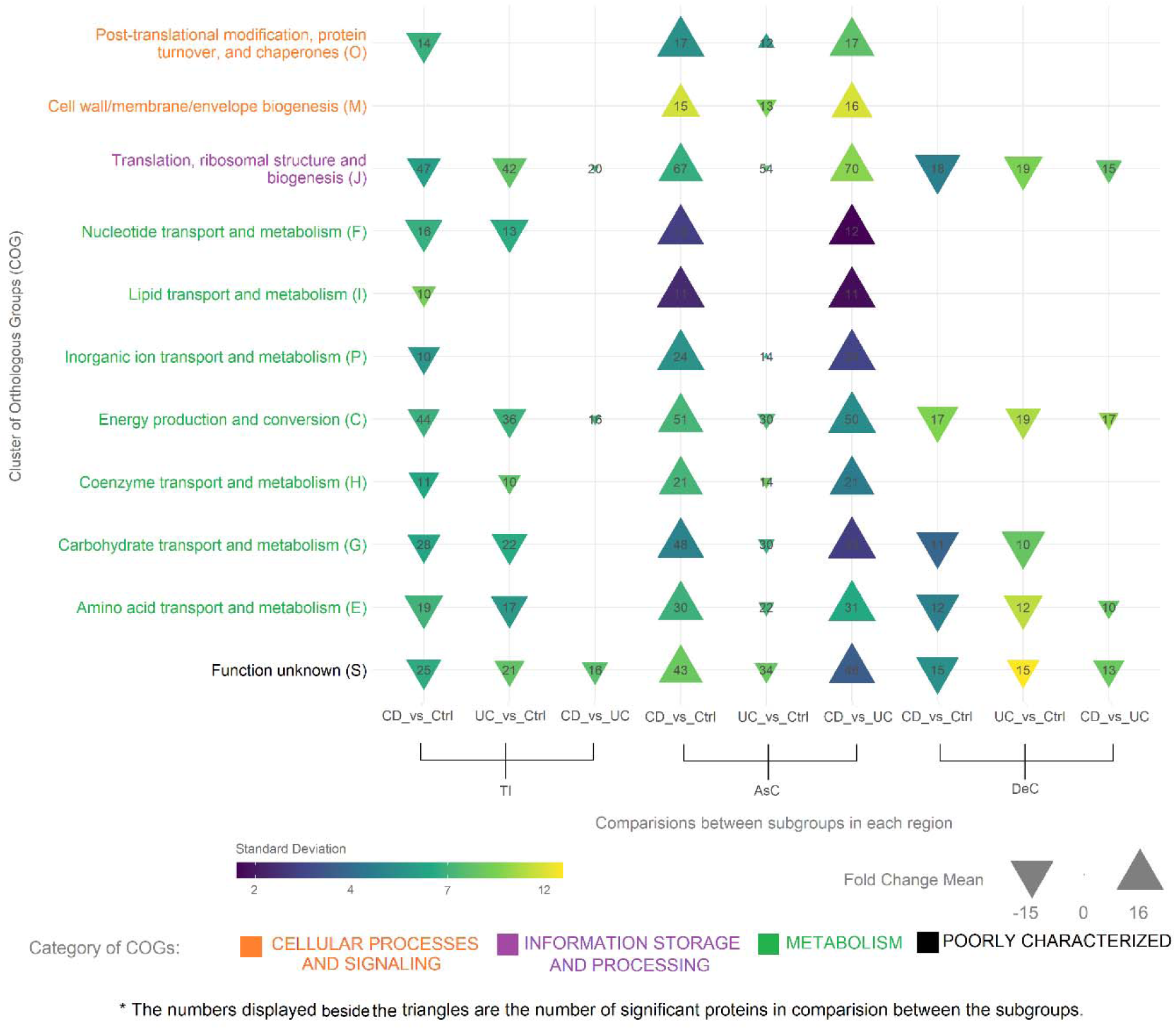
Distinct signature of microbial functionality in the AsC region of CD patients. Cluster of Orthologue Groups (COGs) of significant MPs in the three regions TI, AsC, DeC, among the three subgroups UC, CD, and Ctrl. The size of the triangles correlates with the mean intensity fold change of the significant proteins. Up-pointing and down-pointing triangles demonstrate positive and negative fold change intensity when comparing one subgroup to another. The number of significant proteins for each COG is depicted in the corresponding triangle. Each COGs has at least 5 significant MPs (adjusted *p*-value<=0.05 and absolute fold change>=1.5). TI: Terminal Ileum, AsC: Ascending Colon, DeC: Descending Colon, UC: Ulcerative Colitis, CD: Crohn’s Disease, Ctrl: Healthy control.

In an overall view, it seems that, contrary to the HP-based enrichment analysis (Figure 5), phylogenetic classification of MPs demonstrated that more microbial functionality alterations occurred in the TI compared to the DeC. This is due to the presence of more Q70 proteins in this region (Figure 4). Despite these differences, they showed a relatively similar pattern of functional changes, indicating that the expression levels of these proteins are generally lower in PIBD compared to healthy individuals, confirming dysbiosis as a key feature of the pathogenesis of IBD.

Interestingly, the pattern of changes in the AsC region of CD patients is completely opposite to that of the two other regions. As shown in Figure 6, CD subtype had more pronounced and distinct pattern with positive fold changes relative to healthy control and UC, while fold changes in UC were negative compared to healthy individuals. This indicates that the two subtypes of the disease exhibit nearly opposite changes in terms of microbial protein function compared to healthy individuals. It seems that the highest number of significant proteins between two IBD subtypes (Figure 3E), particularly the presence of a high number of CD-related MPs (Figure 4) in AsC region led to distinguishing this subgroup from the others. These findings are in line with Morgan *et al.* [66, 67], who demonstrated that functional perturbations could be observed in CD with inflammation than UC

In our analysis, functionality changes were based on four fundamental categories: cellular processes and signaling, information storage and processing, metabolism, and a distinctive category embracing functionalities yet to be discerned, labeled as “unknown functionality”. The most enriched COGs were related to increased metabolic activities of the resident microbiome in the AsC of CD patients compared to healthy individuals and UC patients, such as energy production and conversion (C), amino acid transport and metabolism, carbohydrate and energy metabolism, and nucleotide transport and metabolism (F) (Figure 6). Perturbation in the metabolic activity of the gut microbiome has also been reported by Sundqvist et al., reflecting the microbiota’s contribution to the pathogenesis or progression of CD[68].

Another microbial functionality that was significantly altered in IBD patients, particularly in CD ones, compared to healthy individuals belonged to cellular processing and signaling category, such as cell wall/membrane/envelope biogenesis and post-translational modification/protein turnover, and chaperones, suggesting the importance of cell structure integrity and stress response mechanisms in IBD. These processes appear to be more drastically altered in AsC of CD patients compared to UC patients and healthy controls, which could indicate that the microbiome in CD is more stressed or that its microbial community undergoes more substantial changes in composition and function. AsC and TI are mostly involved in more than half of children with CD. It has also been indicated that the bacterial strains are shifted downwards from the AsC to the rectum through the direction of intestinal movement, with the AsC having the highest diversity [69]. These differences might reflect an adaptive response to the chronic inflammation, where the microbial community needs to manage protein folding and maintain cellular integrity.

#### The abundance of bacterial taxa may be associated with a protein that prevents DNA denaturation under extreme conditions

In the taxonomic profiling of significant MPs, we employed eggNOG-mapper to extract information at the family level. In the classification, proteins that were less expressed when comparing one subgroup with another were denoted by the “<” symbol in Figure 7, while overexpressed ones were marked by the “>” symbol. The pie charts in the figure represent the number of significant MPs in each family (details in Supplementary Table S5).

**Figure 7.**
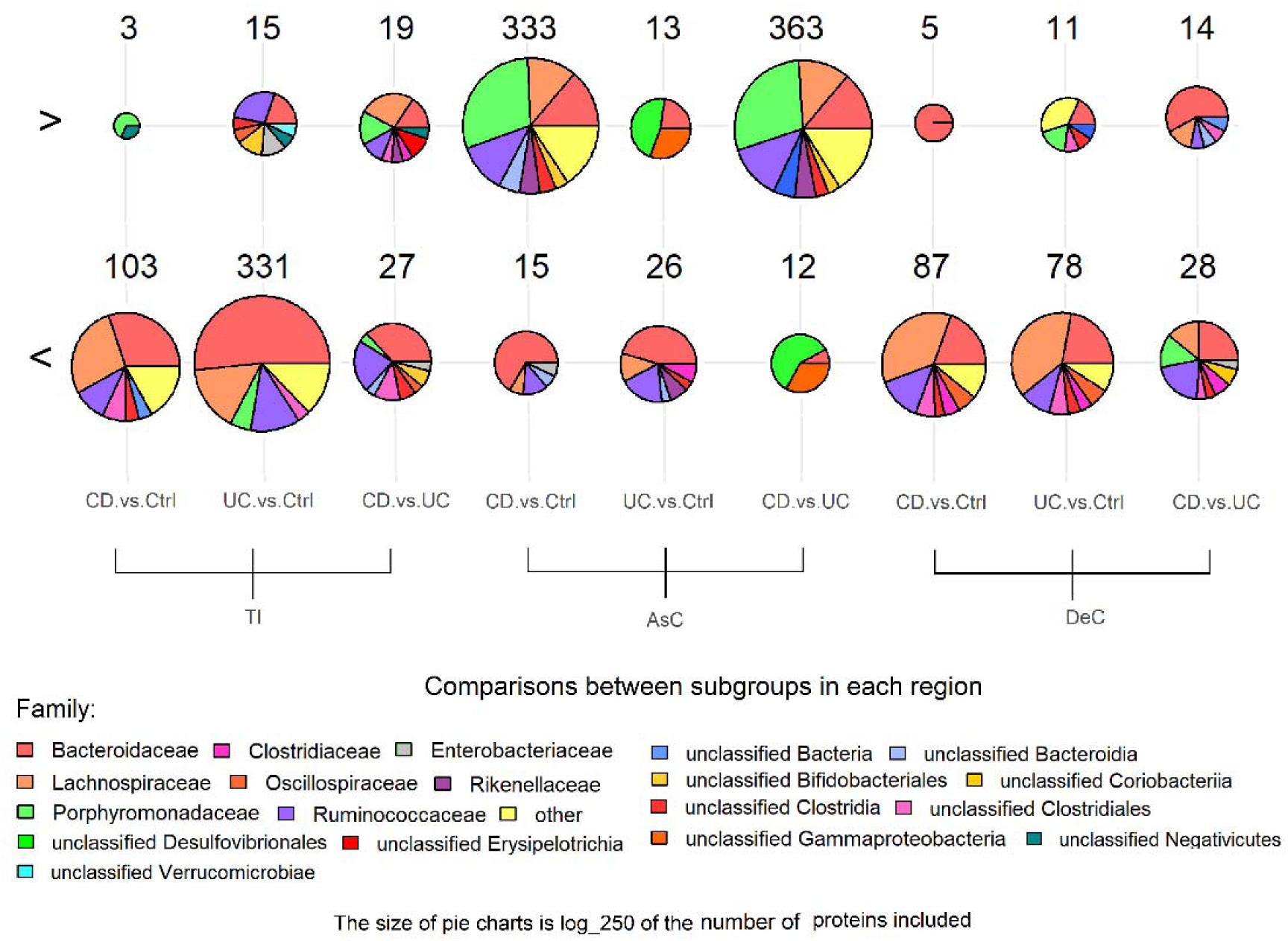
*Bacteroidaceae*, *Porphyromonadaceae,* and *Lachnospiraceae* have the most significant proteins in the TI, AsC, and DeC regions, respectively. Taxonomic profiling of the mucosal luminal interface significant proteins among CD, UC, and control individuals in three regions TI, AsC, and DeC. Down-regulated proteins are demonstrated by the “<” symbol when comparing one subgroup to another, while overexpressed ones are marked by the “>” symbol.

As illustrated in Figure 3, the highest number of significant proteins were found in the AsC region between two IBD subtypes. A noteworthy observation (Figure 7) is that the majority of significant proteins in the AsC region of CD patients compared to UC and healthy individuals belonged to the *Porphyromonadaceae* family, (105 CD-related proteins). To a lesser extent, overexpression of *Porphyromonadaceae* proteins was also observed in UC samples compared to controls. These findings align with previous studies on UC [37], and CD subgroups [39, 70]. *Porphyromonas* ranks among the top ten anaerobic 26 taxa in the microbiome (anaerobiome) of healthy individuals. Bacteria belonging to this family, part of the *Bacteroidota* Phylum, have coccobacilli forms and are strictly anaerobic, non-motile and unable to form spores. In general, scientific literature typically designate *Porphyromonas* species to specific locations in the body, such as *P.gingivalis* and *P.endodontalics* in the mouth, and *P.uenonis* in the gut, and *P.somerae* in the reproductive system [71]. The oral bacteria such as *P.gingivalis* from periodontitis could translocate to the intestine, where they, along with the intestinal microbiome, can lead to dysfunction of intestinal epithelial cell barriers and increase intestinal inflammation. Once oral microbiota pathogens in saliva reach a certain level, they could start colonizing the intestine.

Furthermore, elevated levels of oral bacteria can trigger unusual immune reactions through the movement of T helper 17 cells. *P.gingivalis* oral infection hinders colonic motor functions, indicating that periodontal pathogens can influence the body’s response to IBD. Inflammatory cytokines are expressed and become active in the mouth, and they may travel through the bloodstream to the intestine, affecting IBD. This evidence suggests that periodontitis and colitis may be closely related due to significant changes in the microbiome and *P.gingivalis* plays a role in the development of IBD through the oral–gut disease axis [72, 73].

As mentioned, the majority of significant MPs in the AsC region were CD-related (Figure 4), among which, GL0164822, a *Porphyromonadaceae* protein with 10 unique peptides, functions as a histone-like DNA-binding protein, stabilizing DNA and preventing denaturation under extreme environmental conditions. The histone-like DNA-binding protein associated with the nucleoid is known to be involved in organizing and protecting DNA. These proteins play a role in DNA supercoiling, and compaction, and are also responsible for regulating DNA-associated functions like replication, recombination, repair, and transcription. It has a crucial function in protecting the DNA of bacteria like *H. pylori* from damage caused by oxidative stress and acid stress, which are often faced by the bacterium in the host [74]. Over expression of this protein in IBD, particularly in CD, may indicate an increase in the population of this family or an increase in its activity in the affected area of IBD patients. These findings are also consistent with the number of proteins belonging to this family in the AsC region (Figure 7).

Taxonomic profiling of 351 MPs in the TI region, where the most significant proteins belonged to the Q70 group, revealed that the highest number of down-regulated proteins in UC compared to controls were designated to the *Bacteroidaceae* family. *Bacteroidaceae* are Gram-negative, bile-resistant, motile or nonmotile, and strictly anaerobic bacteria that are the predominant genus in the human gut. They have unique physiological characteristics such as regulation of the intestinal microenvironment, carbohydrate metabolism, and secretion of different metabolite like short-chain fatty acids[75]. Numerous studies underscore the significant potential of *Bacteroides* species in alleviating intestinal inflammation and protecting health. For instance, *B.uniformis* JCM5828 helps reduce colitis by regulating colon NF-κB and MAPK signaling pathways through IL-17 signaling inhibition. *B.uniformis* F18–22 mitigates intestinal inflammation by improving gut dysbiosis. Additionally, *B. fragilis* ZY-312 prevents the adhesion and invasion of intestinal epithelial cells by *Cronobacter sakazakii*, restores intestinal epithelial barrier function, and regulates cytokine expression to reduce necrotizing enterocolitis [76]. Specific species of Bacteroidetes also help regulate inflammation through zwitterionic capsular polysaccharides, which influence T cells by promoting the production of anti-inflammatory interleukin-10-secreting Treg cells [37].

Notably, a significant down-regulated protein in this region of UC patients was GL0091556, which is a histone-like DNA-binding protein, too. This result is consistent with the broader taxonomic profile, indicating a possible reduction in the *Bacteroidaceae* population or decreased expression of related proteins in UC patients. A multiplex gene expression assay of mucosal tissues identified *Bacteroidetes* species’s role in the mucosal immune response. *Bacteroides* species in UC patients showed a significantly reduced relative abundance associated with exacerbation of UC symptoms as these species that attach to the mucosal surface may find it challenging to inhabit the niche of severely damaged mucosa, particularly in the absence of adequate mucin production [37].

Intriguingly, this phenomenon also occurred in the DeC region in which the only significant protein with the function of preventing DNA denaturation belonged to the *Lachnospiraceae* family. This protein was Ctrl-related and had quite low expression levels in both IBD subtypes. This family also has the highest number of proteins with reduced expression in this region, suggesting that this family might have either a decreased population or reduced activity in the patients. The *Lachnospiraceae* are anaerobic, fermentative bacteria involved in strong hydrolyzing activities and producing short-chain fatty acids. In IBD patients’ microbiota, *Lachnospiraceae* have been reported to be reduced in both CD and UC [77, 78].

There is a correlation between the gut microbiota and oxidative stress in both directions. The gut microbiota influences oxidative stress by producing metabolites and antioxidant enzymes and preserving gut homeostasis. On the other hand, oxidative stress could impact the gut microbiota by promoting dysbiosis. The continued presence of ROSs in the intestinal environment could be a causing factor for the development of various diseases, as they contribute to the initiation of persistent and chronic inflammation, immune responses, and DNA damage [79]. Histone-like proteins, small and basic bacterial proteins, are responsible for chromosome dynamics and organization. In addition to their function in DNA compaction, some of them have been shown to have the capacity to protect DNA, thereby maintaining genomic DNA integrity. It is well established that they are important in protecting DNA for adaptation to unfavorable conditions, including different stressors. Histone-like proteins have the capacity to protect DNA from degradation caused by DNAase I and reactive oxygen intermediates. They shield DNA and physically protects the macromolecule by modeling its topology. This activity alters the bacterial transcription profiles, impacting cellular physiology significantly [80, 81].

Considering the COG analysis (Figure 6) and taxonomic enrichment (Figure 7), it becomes evident that distinct signatures of MPs in the AsC region are appropriate for comparing enriched information in IBD subtypes and could be used to distinguish Ctrl from IBD and its subtypes.

The total number of proteins is shown at the top of each pie chart. The sizes of the pies represent the log250 of the number of input proteins. TI: Terminal Ileum, AsC: Ascending Colon, DeC: Descending Colon, UC: Ulcerative Colitis, CD: Crohn’s Disease, Ctrl: Healthy control.

#### A notable difference between the expression pattern of microbial and human features

By applying the LASSO method on the significant proteins, 6 MPs and 6 HPs (denoted by in Figures 2, 3, and 4) were selected from each region (Supplementary Table S6). Interestingly, there were no common proteins among three regions, except for a human protein, which was shared between the TI and AsC regions (Figure 3G).

The boxplots depicted in Figure 8 clearly illustrates a remarkable contrast between the intensity of microbial and human features in the discovery set of 42 patients. Microbial features were mostly absent or newly expressed in the MLI samples of PIBD, while the expression levels of human features typically varied among the subgroups. As shown in Figure 8, the variance of the intensity in the AsC region was lower than in other regions for the human protein features. Furthermore, the expression pattern of the six human features was similar in this region, with their expression level being lower in CD compared to the other subgroups.

**Figure 8.**
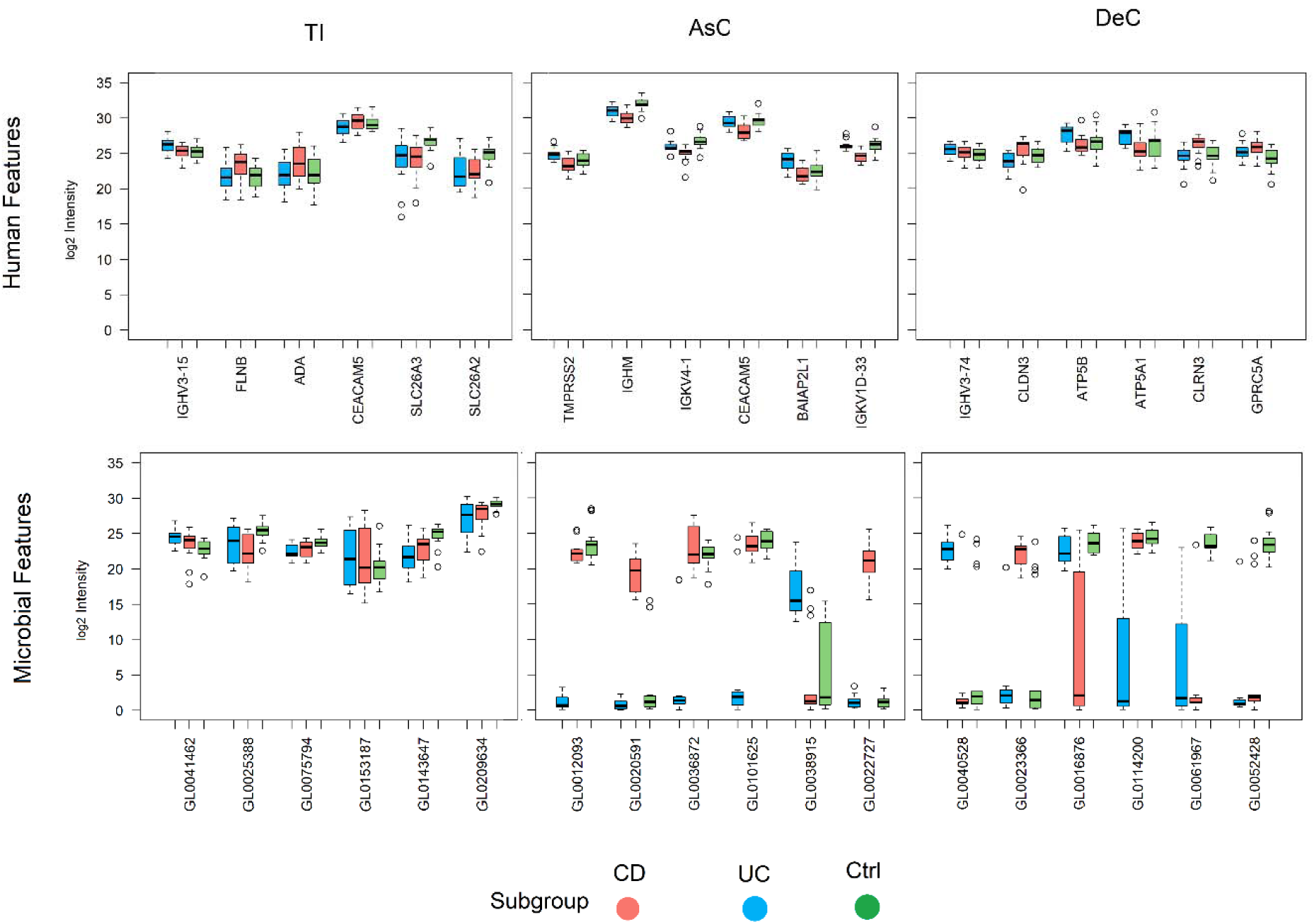
Distinct pattern of expression between microbial and human features. The boxplots indicate log2 intensity of microbial and human features. Microbial features are mostly absent or newly expressed, while the expression levels of human features change among the subgroups. Outliers >1.5 times the IQR are indicated by circles. TI: Terminal Ileum, AsC: Ascending Colon, DeC: Descending Colon, UC: Ulcerative Colitis, CD: Crohn’s Disease, Ctrl: Healthy control.

Intriguingly, all human features in these three regions were from the Q70 protein set, with none belonging to the subgroup-related set. It seems that even when human proteins are subgroup-related, they don’t have sufficient score to be selected as distinguishing features and lack strong discriminatory power. A similar pattern was also observed for microbial proteins in the TI region, where changes in their expression gave them a higher priority for selection as features.

Human features can be categorized based on their functional roles, providing insights into various aspects of IBD pathogenesis. IGHV3-15, IGHM, IGKV4-1, IGKV1D-33, and IGHV3-74 genes are associated with the immunoglobulin heavy and light chains, which are crucial for the adaptive immune response. Differential expression of these immunoglobulin genes may indicate immune dysregulation, a key feature of IBD [50]. Most of them were selected as features in the AsC region and except for IGV3-74 in DeC, the rest were significant between the two IBD subtypes. They contribute to B-cell receptor diversity and antibody production. Transcriptome analysis of colonic biopsies obtained from active UC patients showed overexpression of IgM and IgH heavy chain genes in the active lesions [82]. Likewise, a widespread activation of the humoral immune response has been shown in both CD and UC, as reported by different changes in immunoglobulin subclass production. The interaction of the intestinal immune system with enteric microbes leads to the generation of immunoglobulin (Ig) A and IgG, which are the main antibody isotypes present on the surface of the intestinal mucosa. These antibodies offer defense against infection by attaching to and covering pathogens in the intestinal lumen. In IBD patients, the distinct reactions of the adaptive immune system, which has a greater role in IgG than in IgA production, are altered against these luminal antigens [83].

FLNB, CLDN3, and CEACAM5 are involved in maintaining epithelial cell integrity. FLNB plays a key role in cytoskeletal organization, which is essential for preserving cell structure and function. Disruption of FLNB can affect epithelial barrier integrity, potentially contributing to IBD [50]. CLDN3, a tight junction protein, is also vital for intestinal barrier function. Alterations in CLDN3 can lead to increased intestinal permeability, induced inflammatory response, and gut microbe dysbiosis, common features in IBD [84]. In this study, CLDN3 showed lower expression in the UC subgroup compared to CD and healthy controls. CEACAM5, a cell adhesion molecule, influences epithelial cell interactions and gut barrier function. Carcino Embryogenic Antigen Cellular Adhesion Molecules 5, CEACAM5, was the common feature in the TI and AsC regions, reported before as a dysregulated protein in the biopsy samples of pediatric CD patients and adult UC patients [50, 85]. Interestingly, this gene showed higher expression in the TI region of the CD subgroup, while it had a lower expression in the AsC region of CD patients compared to other subgroups.

Five features, including ADA, SLC26A3, and SLC26A2 in TI as well as ATP5B and ATP5A1 in DeC, are involved in various metabolic processes. ADA is an enzyme crucial for purine metabolism, which affects immune cell function and inflammation. ADA dysregulation in IBD patients has made it an effective biomarker for assessing intestinal inflammation [86]. We showed that ADA expression increased in the TI region of CD patients compared to UC and healthy groups. SLC26A3 and SLC26A2 are solute carriers involved in electrolyte balance and ion transport. Dysregulation of these genes can contribute to diarrhea and electrolyte imbalances observed in IBD. Both Solute Carrier genes showed the same expression pattern among the subgroups, with higher expression in the TI of control samples compared to the two IBD subtypes. SLC26A2 which is involved in maintaining intestinal epithelial barrier integrity [87], has been previously reported as a member of seven-marker genes for differentiating between inflamed and non-inflamed colonic tissue [49]. ATP5B and ATP5A1 are subunits of ATP synthase, which is essential for cellular energy production. Their dysregulation may cause changes in cellular metabolism in inflamed gut tissue [88].

TMPRSS2 which belongs to the serine protease family has been reported to be overexpressed in UC patients [89]. In this line, we showed that this protein is upregulated in UC compared to CD and healthy conditions. GPRC5A is a G protein-coupled receptor whose expression increased in the AsC region of CD and UC patients compared to healthy controls. It is involved in inflammatory signaling and could contribute to the chronic inflammation seen in IBD. Intriguingly, Zhang et al. introduced GPRC5A as a potential biomarker for colon cancer as it promotes tumorigenesis by inducing vanin-1 expression and oxidative stress in colitis associated cancer, a severe complication of IBD [90].

Insulin receptor tyrosine kinase substrate (IRTKS) belongs to IRSp53/MIM family which is involved in the shaping cell membrane protrusions [91]. In fact, this protein contributes to the clustering of actin filaments forming pathogen-driven actin pedestal formation in response to bacterial infections [92, 93]. Furthermore, it has been reported that IRTKS plays a vital role in membrane bending, forming and elongating microvilli on the apical surface of the enterocytes, and regulating microvillus length [94–96]. While direct studies linking IRTKS/BAIAP2L1 to IBD are limited, its known functions in actin cytoskeleton regulation, membrane dynamics, modulating immune responses [97]and cell signaling suggest it as an important player in the pathogenesis of IBD. Our findings showed that IRTKS expression significantly decreased in AsC of CD patients compared to UC ones.

The microbial features highlighted various aspects of bacterial physiology that may contribute to the pathogenesis of PIBD. For instance, proteins such as GL0041462 (MreB, *Verrucomicrobiae*) in TI, which is involved in maintaining bacterial cell shape, and GL0023366 (TonB-dependent receptors, *Bacteroidaceae*) in DeC, which facilitates nutrient acquisition, are essential for bacterial survival and interaction with the host. Although MreB was primarily believed to play a role in determining cell shape, alterations in shape can greatly impact cell physiology and vice versa, making MreB closely linked to various aspects of bacterial growth [98]. The phylum *Verrucomicrobiota* has only a small number of described species. *Akkermansia muciniphila*, a special strain of *Verrucomicrobia*, makes up 1-4% of the overall gut microbiota and lives on the mucosal layer of the intestine by utilizing mucin, a glycoprotein, as its carbon and nitrogen source. *A.muciniphila* regulates the intestinal barrier and immune response by mucosal degradation. Decreased amounts of *A.muciniphila* have been noticed in individuals with IBD, suggesting it could play a role in the progression and onset of IBD, although its effect on IBD is still controversy [99, 100].

The largest number of microorganisms live in the colon and the transit is very slow. As a result, the transport of small molecules across the bacterial outer membrane is a crucial process. Available nutrients are typically large and resistant to breakdown, as fats, proteins, and sugars are digested and absorbed through the small intestine. Fiber, known as dietary polysaccharides, is the primary nutrient source for the colonic microbiota. Furthermore, the abundance of bacterial cells in the colon creates a fierce competition for nutrients. Therefore, the nutrient uptake in *Bacteroides* spp. is mainly facilitated by TonB-dependent transporters rather than diffusion channels [101].

Proteins involved in the metabolic activities of gut bacteria, such as GL0075794 in *Bacteroidaceae*, which is responsible for the conversion of acetate to acetyl phosphate, as well as glyceraldehyde-3-phosphate dehydrogenase (GL0209634 in *Porphyromonadaceae*), which has a function in glycolysis through enzymes, showed greater intensity in the TI of healthy samples, thus playing crucial roles in shaping the gut environment. These processes produce various metabolites that influence host energy metabolism and immune responses. Disruptions in the metabolic pathways of the gastrointestinal microbial community can lead to imbalances in the gut microbiome, which contribute to the pathogenesis of IBD. Indeed, dysbiosis is not only structural changes in the gut microbiota, but can also be related to important dysfunctions in many crucial microbial metabolic activities that may affect the host [67]. Such metabolic dysregulation may result in the production of pro-inflammatory metabolites or the depletion of protective ones, thereby exacerbating gut inflammation.

Additionally, proteins involved in bacterial protein synthesis and DNA regulation, such as ribosomal protein L17 (GL0114200, *Bacteroidaceae*, low intensity in the DeC of UC subtype) and transcription factors with winged helix-turn-helix domains (GL0061967, *Bacteroidaceae*, very low intensity in the DeC of both IBD subtypes) are critical for bacterial growth and adaptation to the gut environment. Ribosomal proteins are widely preserved, abundant and versatile cellular proteins found in all domains of life. The availability of nutrients and the physiological status for bacterial cells in the mammalian gut are constantly changing. During periods of abundance, bacteria like *E. coli* cells in the intestine can have a generation time of 40 minutes, but this can extend to more than 1 day during famine time. As growth rate slows down, both the speed of translation and the amount of ribosomal proteins decrease. In cells that are growing very slowly or are dying, each of these factors may be reduced to almost zero [102]. The winged helix-turn-helix is a common fold in about half of bacterial transcription factors and is the most prevalent effector in all bacterial response regulators, with various DNA regulatory functions [103]. Alterations in these processes could reflect the microbial dysbiosis often observed in IBD patients, where shifts in bacterial populations contribute to disease progression. X-Pro dipeptidyl-peptidase (GL0040528) is a bacterial enzyme whose expression decreased in the DeC of CD patients compared to UC and healthy groups. It is involved in protein degradation and released peptides might influence immune responses in IBD.

GL0012093, GL0022727, and GL0020591 belonging to *Clostridiaceae*, *Lachnoclostridiu*, and *Verrucomicrobiae*, respectively, were among extreme MNAR proteins in the AsC region of CD patients (Figure 2). The functional characteristic of these three features is the putative cell wall binding activity, the glycine cleavage system, and phosphoglycerate kinase, respectively. Bacteria possess different peptidoglycan-binding proteins that play an important role in the cell growth. Cell wall peptidoglycan (PG) growth is supported by dynamic multiprotein complexes that include PG synthases, hydrolases, and cell morphogenesis proteins. Bacteria usually have multiple PG synthases that can polymerize the PG precursor and cross-link the stem peptides. These complexes are positioned and regulated by cytoskeletal elements to form large cell morphogenesis complexes for cell growth (in rod-shaped bacteria) and the divisome for cell division [104]. The overexpression of these proteins could be caused by the increase in bacterial growth and ultimately the increase in the number of bacteria. The previous studies showed an increase in the proportion of *Clostridiaceae* family in CD patients while UC patient’s microbiota has a lower number of this family members [105]. These findings were in line with our results, overexpression of GL0012093 in CD samples compared to its lower expression in UC samples.

Changes in the glycine cleavage system, a multienzyme complex responsible for degrading glycine and linked to energy metabolism [106], further highlights the metabolic divergence between CD and UC. Glycine is a major source of single carbon units for biochemical processes in bacterial cells and is essential for producing various key metabolites, including purines, thymidine, methionine, threonine, and lipids. GL0022727 protein was produced by *Lachnoclostridium*, a member of the *Clostridia* class, in CD samples while it was not detected in UC and control groups. Phosphoglycerate kinase, a key enzyme in glycolysis, is essential for energy production and might contribute to the metabolic shifts in CD. The absence of these proteins in UC may point to distinct microbial profiles and functional pathways that differentiate UC from CD at the molecular level.

#### Microbial features outperform human proteins in diagnostic precision

To minimize the number of markers and propose a panel with the fewest markers while maintaining optimal performance on this dataset, we constructed panels consisting of combinations of 2 to 6 markers for human and microbial proteins in each region. Machine learning was performed using the random forest algorithm for each combination independently, and panels with the highest accuracy on the independent validation set were selected. Supplementary Table 7 provides detailed information on the evaluation of these panels, including F1 scores, the accuracy of the best-performing panel on the training and validation sets, and the number of markers included in each panel.

To explore the feasibility of developing a combined human-microbial marker panel with the minimum number of markers and the highest accuracy, machine learning was applied to all possible combinations of 12 features in each region, ensuring that each combination included at least one human and one microbial protein.

Figure 9 illustrates the ROC curves and AUCs for all panel combinations—human-only, microbial-only, and combined human-microbial markers—in distinguishing each subgroup from the others within each region. The results demonstrate that, regardless of the panel size in the AsC and DeC regions, the overall AUC for microbial features is significantly higher than that for human features. microbial panels exhibited remarkable performance in both the AsC and DeC regions, achieving perfect predictive ability with an AUC of 1 using only two features GL0012093 (*Clostridiaceae*) and GL0022727 (*Lachnoclostridium*) (Figure 8). In contrast, human panel did not reach this level of performance in any region. According to Supplementary Table S6, six different combinations of four microbial features in the AsC region achieved the highest predictive accuracy, distinguishing all subgroups in the validation set with 100% accuracy. Notably, adding even a single human feature to the microbial panel reduced its accuracy compared to the corresponding microbial-only panel.

**Figure 9.**
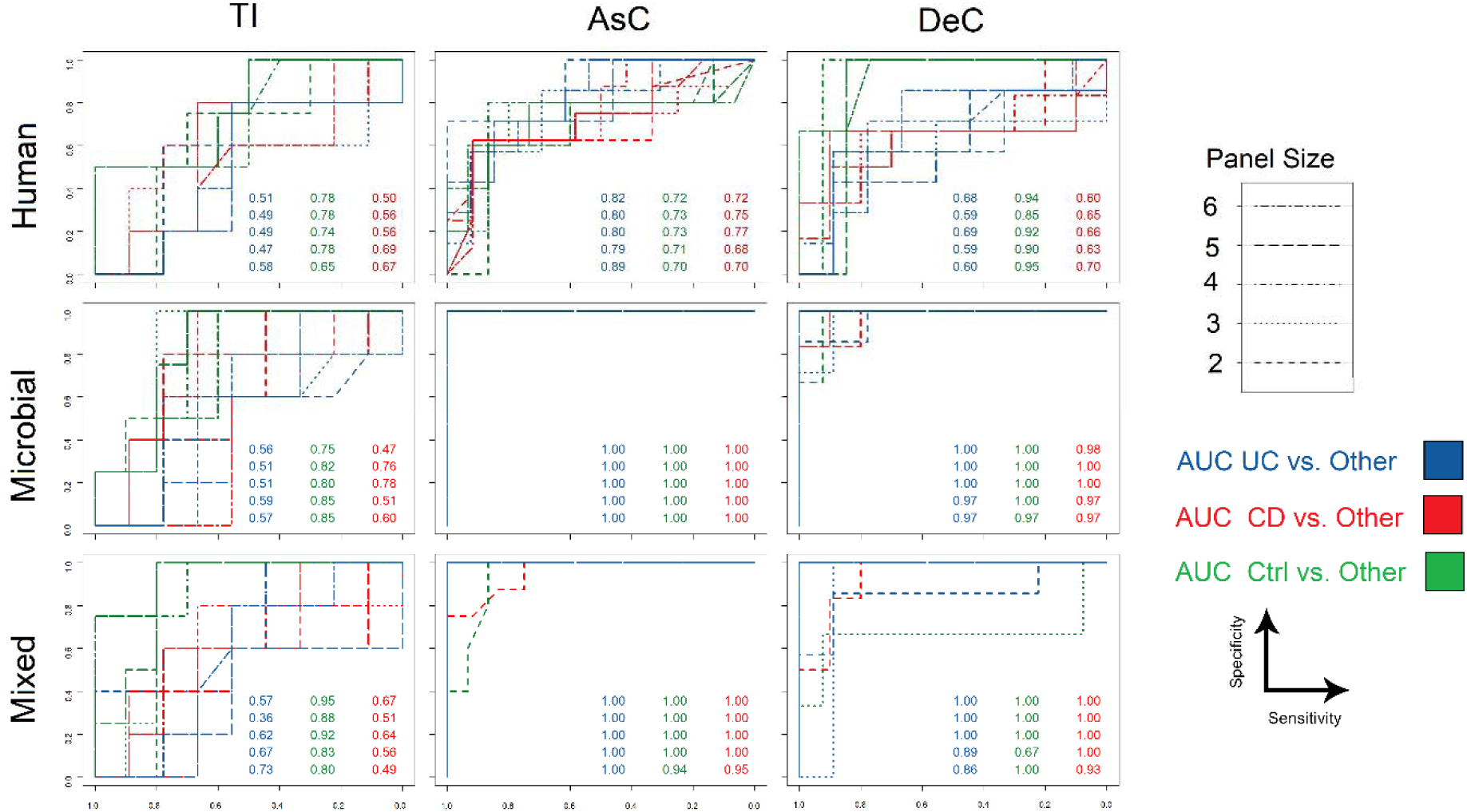
Microbial features of AsC region led to the most accurate biomarker panels for diagnosis Ctrl, UC, and CD pediatric patients. The ROC curves and AUC values for the best combinations of 2 to 6 microbial, human, and combined features on the independent test set in three regions TI, AsC, and DeC, comparing each subgroup against the others. TI: Terminal Ileum, AsC: Ascending Colon, DeC: Descending Colon, UC: Ulcerative Colitis, CD: Crohn’s Disease, Ctrl: Healthy control.

AUC values for panels in the TI region indicated that both human and microbial panels were more effective at distinguishing healthy individuals from IBD patients than differentiating between IBD subtypes. This suggests a lack of significant proteome and metaproteome changes in this region that could support IBD subtype diagnosis. Additionally, while human features demonstrated limited diagnostic accuracy, particularly for distinguishing CD from UC in the DeC region, the inclusion of microbial features significantly improved the accuracy in this region. Although human features exhibited relatively higher diagnostic ability in the AsC region compared to other regions, their performance remained unsatisfactory overall. In contrast, microbial features achieved maximum diagnostic accuracy in the AsC region, even when using only two features. According to Figure 4, feature selection from the subgroup-related proteins of AsC and DeC regions resulted in higher accuracy, whereas those from the TI region which belong to are from the Q70 set demonstrated insufficient diagnostic ability.

Combining human and microbial features slightly improved diagnostic accuracy in the TI and DeC regions. In contrast, the combined human-microbial panels, which included at least one human and one microbial feature, did not enhance performance in the AsC region; instead, they reduced accuracy compared to using microbial features alone. This finding suggests that the predictive power of microbial features surpasses that of human features.

In a previous study [8], researchers proposed a 5-marker based on human proteins obtained from inflamed mucosal biopsy for distinguishing healthy controls from IBD patients. It suggested that the human proteome lack sufficient diagnostic precision for reliably differentiating IBD subtypes.

## 4. Conclusion

Many previous studies on IBD biomarker discovery have focused on host genes and proteins genes. However, numerous studies have also highlighted significant differences in microbiome composition between IBD patients and healthy individuals, particularly in terms of microbial diversity and the relative abundance of certain bacteria. This close relationship between the microbiome and the disease suggests that microbial changes could serve as key diagnostic features. In fact, there are many potential gut microbiome-related biomarkers, including DNA, coding and non-coding RNAs, lipids, metabolites, and proteins, that could be utilized as indicators of disease activity and treatment outcomes in IBD. Alterations in proteomic and metaproteomic profiles provide valuable insights into functional changes during the disease. Therefore, we conducted a comprehensive analysis of protein data obtained from mucosal luminal interface samples of PIBD patients. To explore the diagnostic potential of the microbiome, we separately analyzed human proteome and microbial metaproteome profiles using a dataset derived from both healthy children and those with the disease.

To uncover the latent biomarker potential within the complex gut microbial ecosystem, two classes of proteins were included in the analysis. The first class consisted of proteins with significant expression changes across the disease (Q70). The second group comprised proteins whose presence was entirely disturbed in specific subgroups (subgroup-related). This distinctive type of proteins provided valuable insights for selecting potential biomarkers.

The use of proteomic and metaproteomic data, an appropriate imputation method, and a suitable approach for identifying differentially expressed proteins led to the discovery of PIBD-related pathways, as well as taxonomic and functional insights into the dynamics of the gut microbiota. The feature selection results, along with the machine learning analysis, demonstrated that leveraging the microbial protein profile not only significantly enhanced diagnostic accuracy of the panels but also achieved a 100% accuracy using only two microbial proteins. To our knowledge, this is the first study identifying microbial proteins as biomarkers for IBD diagnosis. The findings clearly showed that the presence or absence of microbial proteins provides better diagnostic ability than relative changes in the human proteome, with purely microbial panels outperforming both human and combined panels.

Remarkably, our findings underscore the ascending colon as the optimal region for mucosal luminal interface sampling and diagnostic biomarker discovery, due to its high diversity in both human and microbial components. This analysis highlights the critical role of region-specific microbial activities and their potential involvement in the pathogenesis and management of IBD. Although the introduced panel demonstrates remarkable performance, we recommend further evaluation in a larger sample-sized dataset, along with laboratory validation, to confirm the generalizability of the suggested biomarker panel and to develop a robust, clinically applicable diagnostic panel for this disease.

## Acknowledgements

We thank Research Institute for Gastroenterology and Liver Diseases, Shahid Beheshti University of Medical Sciences, Tehran, Iran for providing a computational server to perform comprehensive analysis.

## Notes

### Competing Interest Statement

The authors have declared no competing interest.

