## Supplementary figures and images for "Machine Learning-Enhanced Mucosal Luminal Interface Metaproteomics Reveals Diagnostic Potential of Microbial Proteins in Pediatric Inflammatory Bowel Disease"

### suplementary figure 1

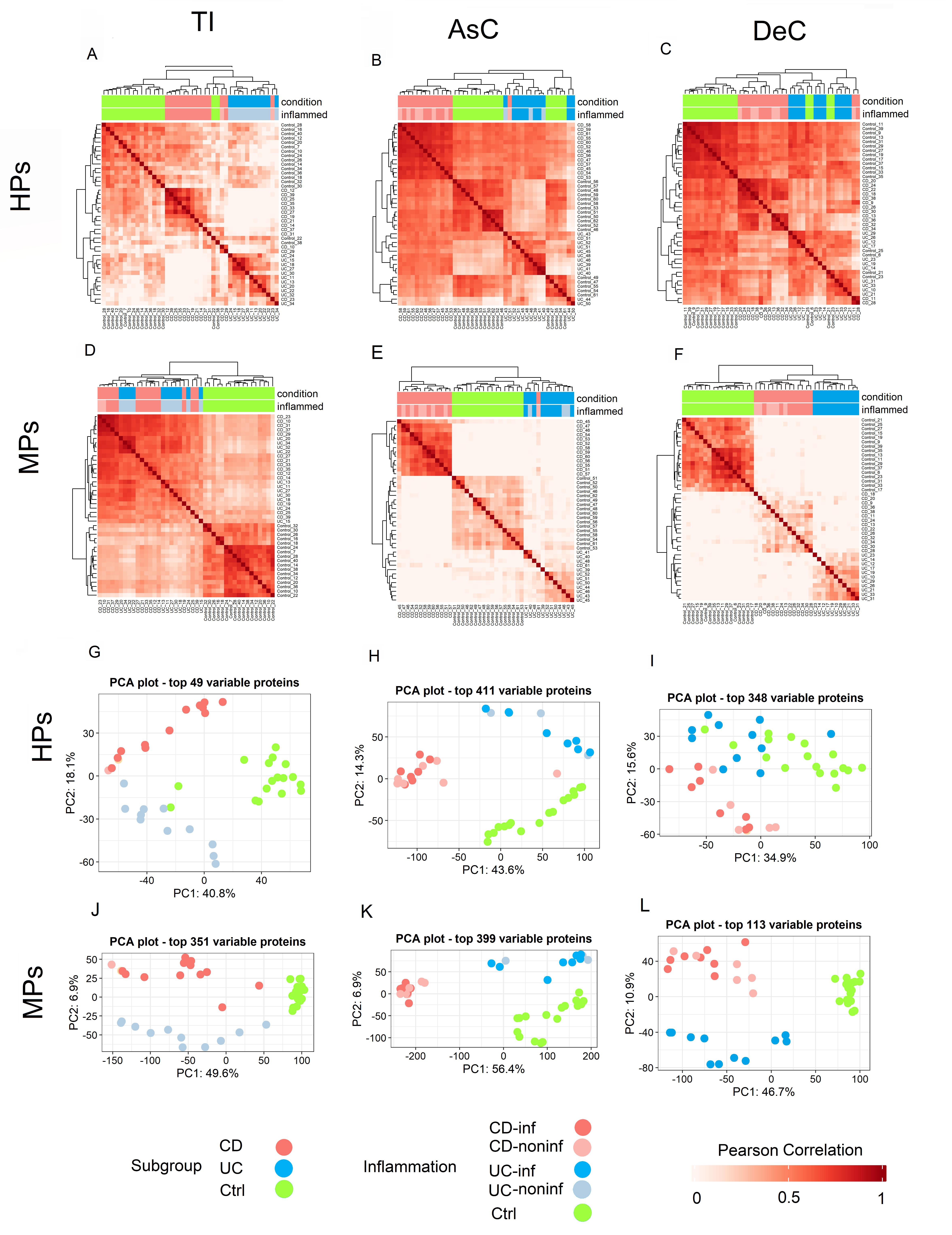
